# Atmospheric CO_2_ availability does not equate to increased nodularin production in diazotrophic cyanobacteria, but does induce varying responses in net photosynthesis and N_2_ fixation rates

**DOI:** 10.1101/203869

**Authors:** Nicola Wannicke, Michelle M. Gehringer

**Affiliations:** Department of Plant Ecology and Systematics, Technical University of Kaiserslautern, Germany.; Leibniz-Institute for Baltic Sea Research, Rostock, Germany; Leibniz-Institute for Plasma Science and Technology, Greifswald, Germany

**Keywords:** nodularin, *Nodularia*, *Nostoc*, climate change, nett photosynthesis, nitrogen fixation

## Abstract

Increasing levels of CO_2_ in the atmosphere are suggested to favour increased incidences of cyanobacterial blooms in water bodies, with a potential concomitant increase in toxin production. As nitrogen fixing cyanobacteria are independent of nitrate and ammonium, this pilot study investigated whether elevated atmospheric CO_2_ levels (eCO_2_), could increase toxin production and net photosynthesis (NP) rates in both terrestrial and aquatic diazotrophic cyanobacteria. Both toxin and non-toxin producing strains of *Nostoc* and *Nodularia* were grown at present atmospheric levels (PAL) of CO_2_ or near future elevated (eCO_2_) and net photosynthesis (NP) determined. Short term responses demonstrated CO_2_ associated increases and decreases in NP, with *N. harveyana* SAG44.85 showing little change in its NP at eCO_2_. Long term responses recorded increases in NP for all species in response to eCO_2_, except for *N. harveyana* on day 7. Nitrogen fixation rates were significantly higher by approx. 10 fold in the aquatic *Nodularia* species compared to the terrestrial *Nostoc* species tested. Moreover, nitrogen fixation rates were not significantly higher at eCO_2_, except for *N. harveyana*. There was no direct correlation between increased nodularin production and eCO_2_ in neither aquatic, nor terrestrial nodularin producing species, however there was a significant correlation between nodularin content and POC:PON ratio for the terrestrial *Nostoc* sp. 73.1 not observed for the aquatic *Nodularia spumigena* CCY9414.

## Introduction

Cyanobacteria in their role as primary producers form an essential part of the global C and N cycles, both in terrestrial and aquatic environments. The process of oxidative photosynthesis, whereby CO_2_ fixation is powered through the splitting water, energised by sunlight, is thought to have evolved during the Archean era when there was no free oxygen in the Earth’s atmosphere (Lyons *et al.*, 2014). Cyanobacteria are attributed with oxygenation of the Earth’s atmosphere around 2.33 Ga during a period known as the great Oxygenation Event (GOE), marking the end of the Archean era (Lyons *et al.*, 2014). Phylogenetic studies indicate that all cyanobacterial photosystem II reaction centre D1 proteins evolved before the last common ancestor of cyanobacteria (Cardona *et al.*, 2015), currently positioned at approximately 2.7 Ga (Sanchez-Baracaldo, 2015), indicating that photochemical splitting of water was possible before the GOE. Levels of atmospheric CO_2_ are thought to have been in the range of 5 − 10× present atmospheric levels (PAL) during the late Archean meaning that oxidative photosynthesis evolved under atmospheric CO_2_ levels similar to those proposed under worst case scenario climate change models ~ 2000 ppm. Dating the origin of N_2_ fixation is more complicated but the molybdenum based nitrogenase was recently placed in the last universal common ancestor (LUCA), before cyanobacteria evolved (Weiss *et al.*, 2016).

The enzyme catalysing CO_2_ fixation in Cyanobacteria and modern-day C3 plants is ribulose-1,5-bisphosphate carboxylase/oxygenase (Rubisco), thought to be the most abundant enzyme on Earth. Rubisco binds CO_2_ and generates 2 molecules of 3-phosphoglygerate (3PGA) which is further processed in the Calvin-Benson-Bassham (CBB) cycle to produce ribulose-1,5-biphosphate, Rubisco’s CO_2_ acceptor molecule, and glutamate. In order to reduce undesirable oxygenase activity, cyanobacteria have evolved the carbon concentrating mechanism (CCM) to increase the effective concentration of CO_2_ around the Rubisco active site by up to 1000-fold (Price, 2011). The increased growth rates observed for *Trichodesmium* cultures growing under 900 ppm CO_2_ were ascribed to down regulation of the CCM, thereby reducing the energy demands on the cell (Kranz *et al.*, 2011). In contrast, biological soil crusts showed a decrease in cyanobacterial abundance when grown under eCO_2_ for 10 years, suggesting a negative impact of climate change on arid soil crusts (Steven *et al.*, 2012). The ability of cyanobacterial soil crusts to increase NP under eCO_2_ exposure is dependent on water availability (Lane *et al.*, 2013).

Approximately a third of all anthropogenic CO_2_ released dissolves in the oceans, reducing the pH by increasing the partial pressure of CO_2_. Oxidative photosynthetic organisms that rely on the construction of a carbon concentrating mechanism (CCM) are thought to be sensitive to changes in pCO_2_ (Shi *et al.*, 2012). Decreasing the medium pH reduced N_2_ fixation rates in *Trichodesmium*, an important cyanobacterial, non-heterocystous marine diazotroph, under Fe- limiting conditions. The reduced N_2_ fixation rates corresponded to reduced nitrogenase efficiency at lower pH. Exposing cultures of *Nostoc muscorum* to raised HCO_3_^−^ concentrations under diazotrophic conditions resulted in enhanced growth, O_2_ and pigment production and nitrogenase activities (Bhargava *et al.*, 2013). *Nodularia spumigena* KAC12 grown at eCO_2_ of 960 ppm showed increased photochemical yield after 5 days (Karlberg & Wulff, 2013), suggesting higher potential NP rates. *Nodularia spumigena* CCY9414 grown under elevated CO_2_ conditions (548 ppm) exhibited increased C fixation rates compared to control cultures, with increased carbon to nitrogen (POC:PON) and nitrogen to phosphate ratios recorded (Wannicke *et al.*, 2012). Higher carbon to nutrient ratios were also observed in *Synechocystis* PCC6803 cultures grown at eCO_2_ (Verschoor *et al.*, 2013). Only a slight increase was observed in C:N ratios in 3 strains of cyanobacteria grown at eCO2 in continuous culture in bubble reactors, namely *Cyanothece* ATCC51142, *Nodularia spumigena* IOW-2000/1 and *Calothrix rhizosoeniae* SC01 (Eichner *et al.*, 2014). This excellent study emphasised the need to generate more data on N_2_ fixation responses in heterocystous cyanobacteria in response to eCO_2_ levels, and highlighted the diversity of responses in marine species to eCO_2_ of approx. 900 ppm.

Given their evolutionary history under raised CO_2_ levels, researchers (Gehringer & Wannicke, 2014, Sandrini *et al.*, 2016, Visser *et al.*, 2016) have voiced concerns for increased bloom occurrence and toxin production under the elevated levels of CO_2_ proposed by current climate change scenarios. Particularly toxin producing strains capable of fixing their own N_2_ from the atmosphere offer the biggest threat to human safety, as they would have little nutrient limitations restricting growth (Wannicke *et al.*, 2012). The most commonly occurring cyanobacterial toxins are microcystin and nodularin, both strong protein phosphatase inhibitors, capable of inducing extensive hepatocellular bleeding and collapse in exposed individuals and animals (Gehringer, 2004). The levels of toxin production within cyanobacterial blooms is largely determined by several abiotic factors such as light intensity and quality, pH and nutrient availability (Reviewed by Neilan *et al.*, 2013; Gehringer & Wannicke, 2014; Visser *et al.*, 2016). Raised temperatures and elevated CO_2_ levels in the range of those proposed under climate change, are linked to increased primary production (Paerl & Huisman, 2009) and toxin production by cyanobacteria (El-Shehawy *et al.*, 2012, Kleinteich *et al.*, 2012). Increased production of the secondary metabolite microcystin is linked to maintaining the C:N balance in the cell (Downing *et al.*, 2005) particularly when N uptake exceeds the relative growth rate. Elevated CO_2_ levels have the capacity to significantly affect the community composition and toxicity of *Microcystis* blooms (Liu *et al.*, 2016; Van De Waal *et al.*, 2011). Microcystin synthesis requires active photosynthesis (Sevilla *et al.*, 2012) and is regulated by the global N uptake regulator, NtcA, supporting the proposed importance of the C:N balance on toxin production. This agrees with observed anthropogenically alterations in the N/P ratio resulting in the appearance of cyanobacterial blooms (Beversdorf *et al.*, 2013) and increased toxin production (Horst *et al.*, 2014). N limitation is thought to induce a shift to N_2_ fixing, diazotrophic cyanobacteria, thereby increasing organic N availability and a subsequent increase in toxin production (Posch *et al.*, 2012, Gehringer & Wannicke, 2014).

Dense cyanobacterial blooms require excessive CO_2_ to support their continued growth (Paerl & Huisman, 2009) with CO_2_ availability often limiting bloom growth, a restriction that could be removed under increased atmospheric CO_2_ levels. Only aquatic cyanobacterial strains carrying the high flux, low affinity HCO_3_^−^ receptor were able to benefit from eCO_2_ levels and increase their growth rates (Sandrini *et al.*, 2015, Sandrini *et al.*, 2016, Visser *et al.*, 2016). At the time of our review (Gehringer & Wannicke, 2014) most eCO_2_ studies had focused on the freshwater unicellular non-diazotrophic *Microcystis aeruginosa* species, with only a couple investigating diazotrophs. Subsequently, we decided to conduct a pilot study on diazotrophic cyanobacterial species from both terrestrial and aquatic environments to investigate the effect of eCO_2_ levels on NP, N_2_ fixation rates and toxin production.

## Materials and methods

### Cultures and culture conditions

Terrestrial diazotrophs *Nostoc* spp. 73.1, 40.5, 65.1 and C1.8 were grown on BG11 medium or nitrogen free medium, BG11_0_ medium with ferric ammonium citrate replaced with ferric citrate (Gehringer *et al.*, 2010). The aquatic diazotrophic *Nodularia* species *Nodularia spumigena* CCY9414 (Culture Collection Yerseke) and the benthic *Nodularia harveyana* SAG44.85 (Culture Collection of Algae, SAG, Georg August University, Göttingen) were cultivated in nitrogen free F/2 medium (UTEX, Austin) containing vitamins, with *N. harveyana* requiring the addition of 5 ml l^−1^ of soil extract. *Nostoc* sp. 73.1, *Nostoc* sp. 65.1 (Gehringer *et al.*, 2012) and *Nodularia spumigena* CCY9414 (Voss *et al.*, 2013) produce nodularin.

The experimental flow is illustrated in Supplementary Figure 1. Fifty ml of stationary phase cultures were inoculated into 150 ml of the appropriate media in a glass rim culture flask (Duran, 45 mm dia.) and placed at the control or experimental conditions for 14 days (acclimation phase). The inoculum cultures were then diluted 1:1 with fresh medium and divided into two ventilated T_175_ polystyrene cell culture flasks (Greiner) and laid flat to ensure an even light exposure and maximise gas exchange. Doing the experiments in duplicate permitted the investigation of more diazotrophic species and conditions during this pilot study. Experimental cultures were exposed to eCO_2_ of 2000 ppm, 10:14 hr light:dark cycle, 22 °C / 18 °C light:dark, 60% humidity and 130 μmol photons m^−2^ s^−1^ (Percival chamber). Control cultures were exposed to CO_2_ at PAL (440 ppm in Kaiserslautern, Germany), 10:14 hr light:dark cycle, 22 °C, 60% humidity and 130 μmol photons m^−2^ s^−1^ (Osram L58 daylight bulb, W77).

### Gas exchange measurements, chlorophyll and toxin extraction

Culture material was removed by pipetting under sterile conditions in a clean bench in PAL CO_2_ conditions on days 7 (exponential growth phase) and 14 (stationary phase). Harvested bacteria were filtered onto a 3 μM SSWP glass fibre filter (Millipore), placed onto an appropriate, moist agar plate and incubated under experimental conditions until net photosynthesis (NP) determinations (between 1 to 4 hours after sampling) by means of CO_2_ gas exchange (GFS 3000, WALZ, Effeltrich, Germany). Bacterial covered filters were placed in the sample cuvette and NP determined at 80% humidity first at PAL for control cultures (C_amb_) and then 1000 ppm CO_2_ (C_high_). Cultures treated at eCO_2_ levels were first measured at 1000 ppm CO_2_ (T_high_) and then PAL (T_amb_). The respiration rate was determined for each filter after 5 min dark incubation at the start and end of the measuring period to ensure the cultures were not stressed. CO_2_ assimilation rates were determined at 50, 130 and 500 (light saturation point) μmol photons m^−2^ s^−1^, and expressed per μg chlorophyll *a*. Responses of net photosynthesis to eCO_2_ levels have been split into 2 categories (Suppl. Fig. 1)– short term responses to immediate changes in CO_2_ availability for days 7 and 14 (C_high_ versus C_amb_) and (T_high_ versus T_amb_), as well as long term responses for days 7 and 14 (C_amb_ versus T_high_) measured after 14 days growth under elevated CO_2_. Data for one replicate of *Nostoc* 40.5 grown at eCO_2_ on day 14 indicated that the sample was stressed and it was also excluded from the analysis (T_amb_ & T_high_), while the readings for C_high_ for *N. spumigena* CCY9414 on day 7 were erratic and excluded from our calculations.

A combined protocol for chlorophyll *a* determination and toxin extraction from the bacterial filter discs was employed. After NP measurements, each filter was placed in a 2 ml centrifuge tube containing 100 mg beads (0.1 mm zirconia silica beads, BioSpec) to which 1.5 ml 90% HPLC grade methanol was added (Meeks & Castenholz, 1971). The samples were bead-beated (Retch) for 1 min at 30 beats per min and incubated at room temperature in the dark overnight. Samples were subsequently centrifuged at 10 000 rpm for 5 minutes at 20°C and the OD_665_ of the SNF determined (Lambda 35 UV/VIS spectrometer, Perkin-Elmer). Chlorophyll *a* content was calculated using the equation: Chl *a* μg ml^−1^ = OD_665_ × 12.7 (Meeks & Castenholz, 1971). The SNF was then stored in a new 2 ml centrifuge tube at −20°C. The above protocol was repeated with 1 ml of 70% HPLC grade methanol to complete toxin extraction (Gehringer *et al.*, 2012). Two hundred microliters of a 1:1 mixture of the 90% and 70% methanol extracts were used in a competitive ELISA assay (Beacon Analytical Systems Inc., Portland, ME, USA) following the manufacturer’s instructions. The amount of toxin extracted from each bacterial filter was calculated from the standard curve and expressed as ng nodularin μg Chl a^−1^.

### Stable isotope and N_2_ fixation

Stable N isotope ratios (δ^15^N-PON) as well as PON and POC concentrations were measured for both *Nodularia* samples and *Nostoc* sp. 73.1 grown in N-free medium on day 14 by means of flash combustion in a Carlo Erba EA 1108 at 1020 °C in a Thermo Finnigan Delta S mass-spectrometer. Filters containing culture samples were trimmed, sectioned, then loaded into tin capsules and palletised for isotopic analysis. The stable N isotope ratios measured for each sample were corrected against the values obtained from standards with defined nitrogen and carbon isotopic compositions (IAEA-N1, IAEA-N2, NBS 22, and IAEA-CH-6) by mass balance. Values are reported relative to atmospheric N_2_ (δ^15^N). The analytical precision for both stable isotope ratios was ±0.2%_0_. Calibration material for N analysis was acetanilide (Merck). N_2_ fixation activity was measured by incubating cultures with bubble addition of ^15^N-N_2_ enriched gas for 24 hours, guaranteeing sufficient dissolution of the ^15^N gas in the incubation bottle (Mohr *et al.*, 2010). Tracer incubations were terminated by gentle vacuum filtration (<200 mbar) of the culture material through pre-combusted GF/F filters (Whatman) that were then dried at 60°C, analysed and the N_2_ fixation rates calculated (Montoya *et al.*, 1996).

### Carbonate Chemistry

The pH was determined on day 14 from sample filtrates using an electrode (Radiometer analytical PHM210, France) calibrated with a three point calibration using NBS (National Bureau of Standards) buffers giving values of pH relative to the NBS scale. Total alkalinity (A_T_) was determined using the colorimetric SOMMA system according to Johnson et al. (1993). The system was calibrated with carbon reference material provided by A. Dickson (University of California, San Diego) and yielded a precision of about ±2 μmol kg^−1^. Total carbon (C_T_) and pCO_2_ in the media were calculated using CO2SYS (Lewis *et al.*, 1998).

### Statistical analysis

Statistical analyses were done by using repeated measures ANOVA to determine the CO_2_ treatment effect and the effect of sampling time on the sort-term response of C assimilation. Furthermore, we applied Spearman's rank correlation to test for dependencies in between experimental parameters. Additionally, Mann-Whitney Rank Sum Test was applied when comparing NP and respiration rates between eCO_2_ treatments and PAL controls. Prior to ANOVA and correlation analysis, data were tested for normality and homogeneity of variances using Wilk-Shapiro and Levene’s tests. All analyses were performed using the software SPSS 22.0.

## Results

### Carbonate Chemistry

Carbonate chemistry confirmed that our experimental application of a continuous gas supply ensured culture growth at elevated CO_2_ levels. The *p*CO2 in the media of cultures grown at 2000 ppm was determined to be in the range of 1987 μatm for BG11_0_ and 1701 μatm for F/2 medium (Suppl. Table 1).

### Net photosynthesis

#### Growth phase response

When considering net photosynthesis rates measured under the growth conditions of PAL and eCO_2_, only *N. harveyana* SAG44.85 grown at PAL displayed an incubation time effect with net photosynthesis higher on day 7 (exponential phase), compared to early stationary phase on day 14. Differences between the two sampling days in all other cultures were observed, nevertheless, a repeated measure ANOVA with time as a predictor for combinations of growth and treatment CO_2_ revealed a significant time effect in *N.spumigena* at PAL (higher rates at day 14) and for *Nostoc* sp. C1.8 (higher at day 14) and *Nostoc* sp. 73.1 (higher at day 7) (with DIN addition) at eCO_2_ (Suppl. Table 2).

During exponential growth at day 7, nitrogen availability (addition of nitrate) seemed to have no effect on NP in *Nostoc* C1.8 and *Nostoc* sp. 73.1 at PAL growth conditions. At eCO_2_, NP in *Nostoc* C1.8 seemed unaffected by N availability, but appeared higher in *Nostoc* sp. 73.1 cultures with DIN addition. No difference in NP was detected between the *Nodularia* species tested at PAL.

During stationary growth (day 14), N availability had no effect on NP in *Nostoc* C1.8 at PAL, whereas at eCO_2_ higher NP was detected in cultures with DIN addition. In *Nostoc* sp. 73.1 no effect of N availability on NP was shown, neither at PAL nor eCO_2_. *Nodularia* species did not display different NP rates at PAL, while at eCO_2_ elevated NP was observed in *N. spumigena* compared to *N. harveyana*.

**Figure 1:**
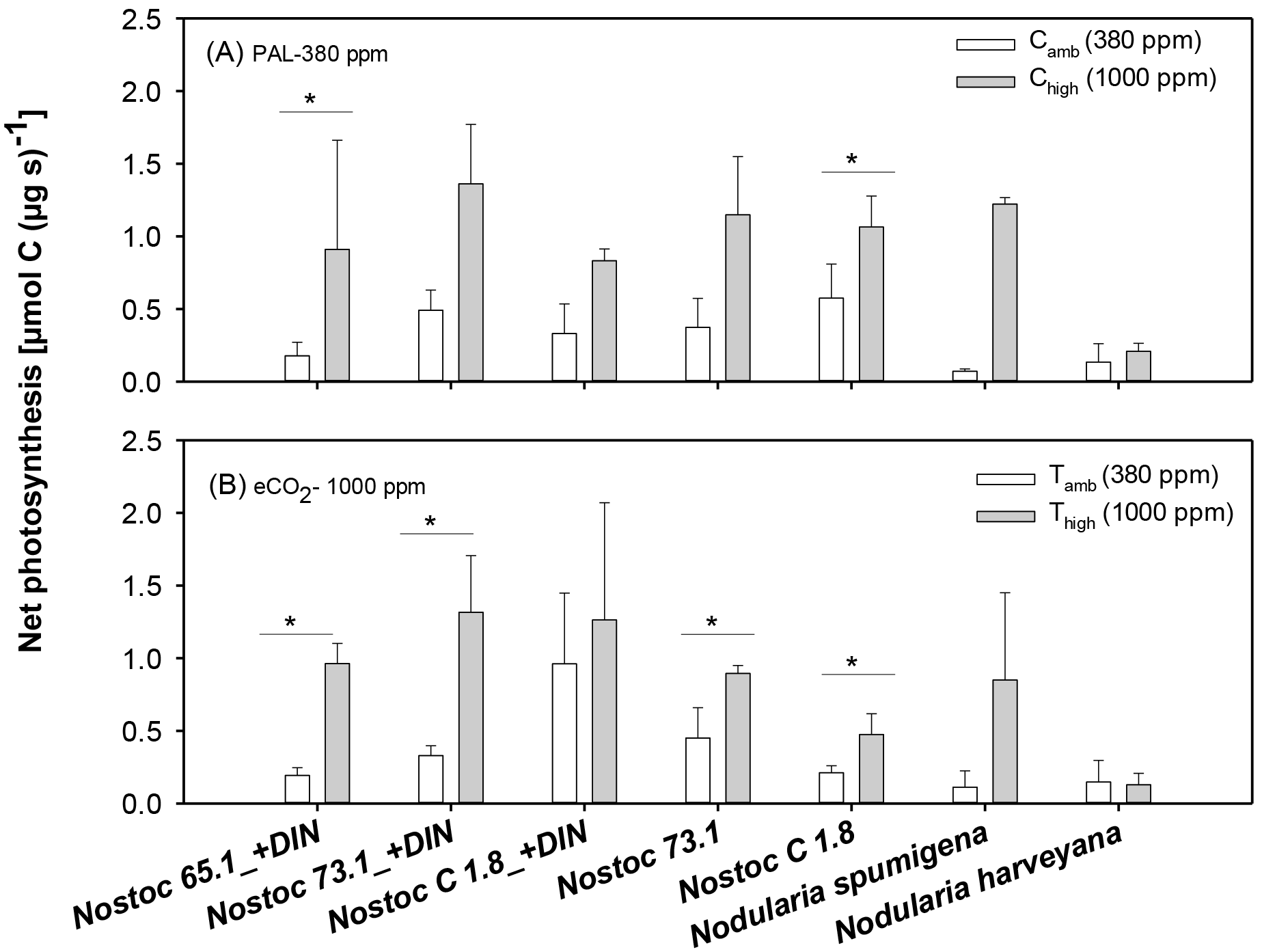
**Short term responses** of net photosynthesis in control cultures (A) grown at PAL (380 ppm) and the treatment (B) grown at eCO_2_ (2000 ppm) with NP (C_amb_ & T_amb_) measured under PAL (380 ppm, white bars) and eCO_2_ (1000 ppm, grey bars) conditions (C_high_ & T_high_). Bars represent mean and standard deviation of n=4 (day 7+day 14). Significant differences between measurements are indicated by * (p≤0.05) according to repeated -measure ANOVA and **(p≤0.005) (Suppl. Table 2).

### Short term responses to eCO2

All control cultures grown and measured at PAL (C_amb_) increased their mean net photosynthesis when presented with 1000 ppm CO_2_ in the sample cuvette (C_high_) (Fig. 1A). This short term response is strain dependent and only statistically significant in *Nostoc* 65.1 (with DIN present) and *Nostoc* sp. C1.8 (without DIN) (Suppl. Table 2). Calculating the logarithmic response ratio of net photosynthesis (NP at C_high_/C_amb_, Supp. Fig. 2A) again reveals an instant increase of eCO_2_ on net photosynthesis in almost all cultures and at both time points, except in *N. harveyana* SAG44.85 at day 7 and most pronounced in *N. spumigena*. In eCO_2_ cultures grown at 2000 ppm, net photosynthesis was significantly elevated at high CO_2_ short- term treatment in *Nostoc* spp. 65.1 and 73.1 (with DIN), as well as in *Nostoc* spp. 73.1 and C1.8 (without DIN) (Fig. 1B, Suppl. Table 2). The logarithmic response ratio of NP also displays a stimulating effect of eCO_2_ on net photosynthesis in all cultures, except *N. harveyana* SAG44.85 (day 7) and was most pronounced in *N. spumigena* CCY9414.

### Long term responses to eCO2

Increased net photosyntheis rates were observed in all *Nostoc* strains grown under eCO_2_ and in the presence of DIN when compared to the PAL control cultures at both logarithmic (day 7) and early stationary phases (day 14), as illustrated using the ln response ratio (Fig. 2). *Nostoc* strains grown without addition of DIN displayed no significant effect of eCO_2_ on net photosynthesis. The effect in the *Nodularia* species was different for the two sampling intervals, displaying a stimulation of eCO_2_ in the exponential growth phase (day 7) in *N. spumigena* CCY9414 and no effect in the stationary growth phase (day 14). In *N. harveyana* SAG44.85, eCO_2_ showed no effect on net photosynthesis at day 7 and a non-significant negative effect at day 14 (Fig. 2).

**Figure 2:**
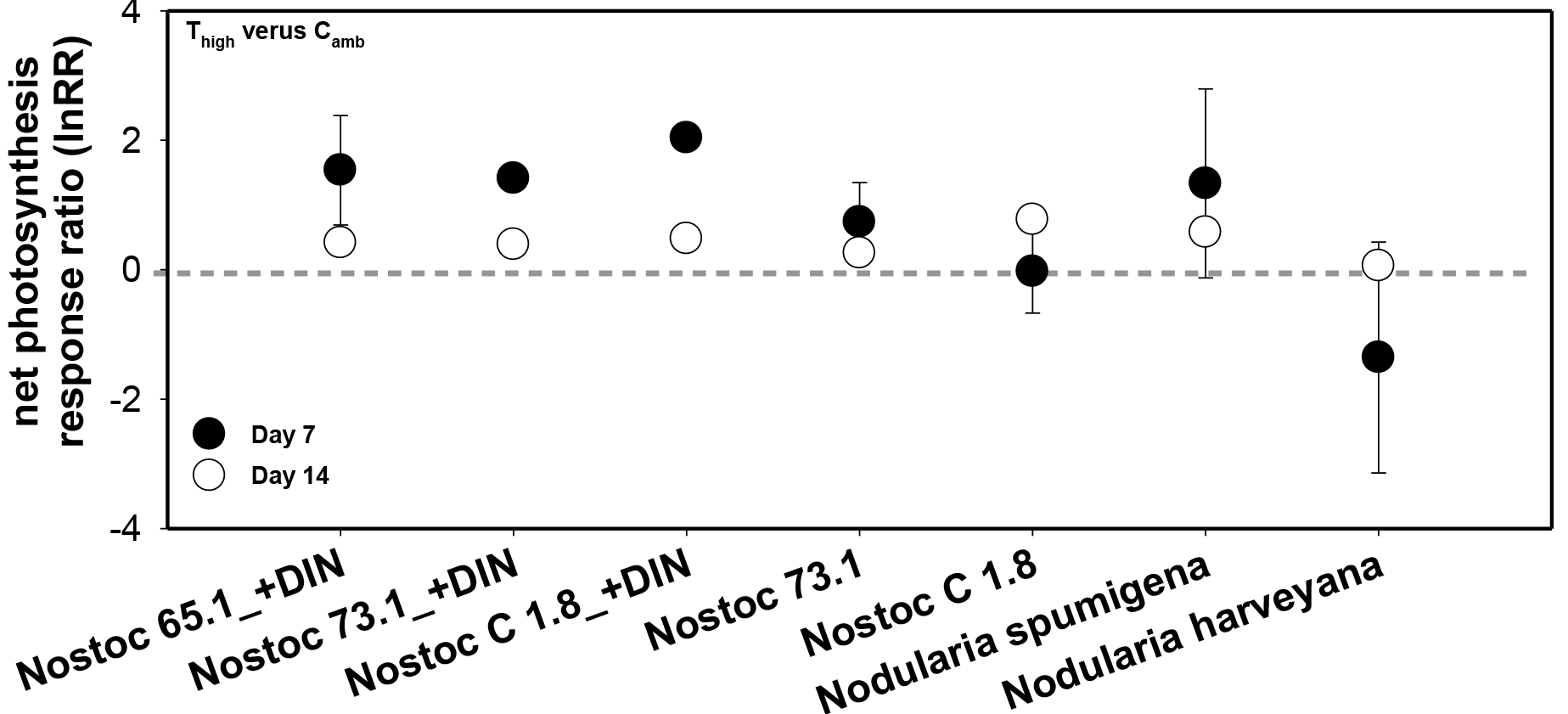
**Long term response** of net photosynthesis (lnRR =natural-log transformed) for the different species tested. LnRR was calculated by dividing net photosynthesis (NP) rate of culture grown at eCO_2_ conditions and NP measured at eCO2 (T_high_) by those of cultures grown at PAL and NP determined at PAL (C_ambient_) conditions. Data is presented as means and 95% confidence interval (CI) of day 7 and day 14, n=2 respectively. The horizontal grey line indicates lack of response to the CO_2_ treatment (i.e. lnRR = 0). If the CI crossed the 0 response line, the effect of eCO_2_ is considered as non-significant. Mean and CI >0 indicate a stimulation by eCO2. Mean and CI <0 indicate a negative effect of eCO2.

### Respiration

No significant trend in respiration was detectable on the days sampled, except for *N. spumigena* CCY9414 grown at PAL, where respiration appeared elevated. This increase was not statistically significant (Mann-Whitney Rank Sum Test, t=15, p=0.1, n=4) at the eCO_2_ treatment relative to PAL (Suppl. Figure 2). There was however a correlation of dark respiration with chlorophyll *a* content and N_2_ fixation rates (positive) (Suppl. Table 3).

### N_2_ fixation

N_2_ fixation rates in both *Nodularia* species were elevated by approximately tenfold when compared to rates determined for *Nostoc* sp. 73.1 (Fig.3). Moreover, N_2_ rates were much higher for *N. harveyana* SAG 44.85 under eCO_2_ growth conditions when compared to the control cultures. There was a trend towards elevated N_2_ fixation rates in *N. spumigena* as well, but this effect was not statistically significant, as well as the reduced N_2_ fixation of *Nostoc* sp. 73.1 at eCO_2_ conditions compared to the control PAL. N_2_ fixation correlated positively with chlorophyll *a* and PON content and negatively with net photosynthesis (Suppl Table 3).

**Figure. 3:**
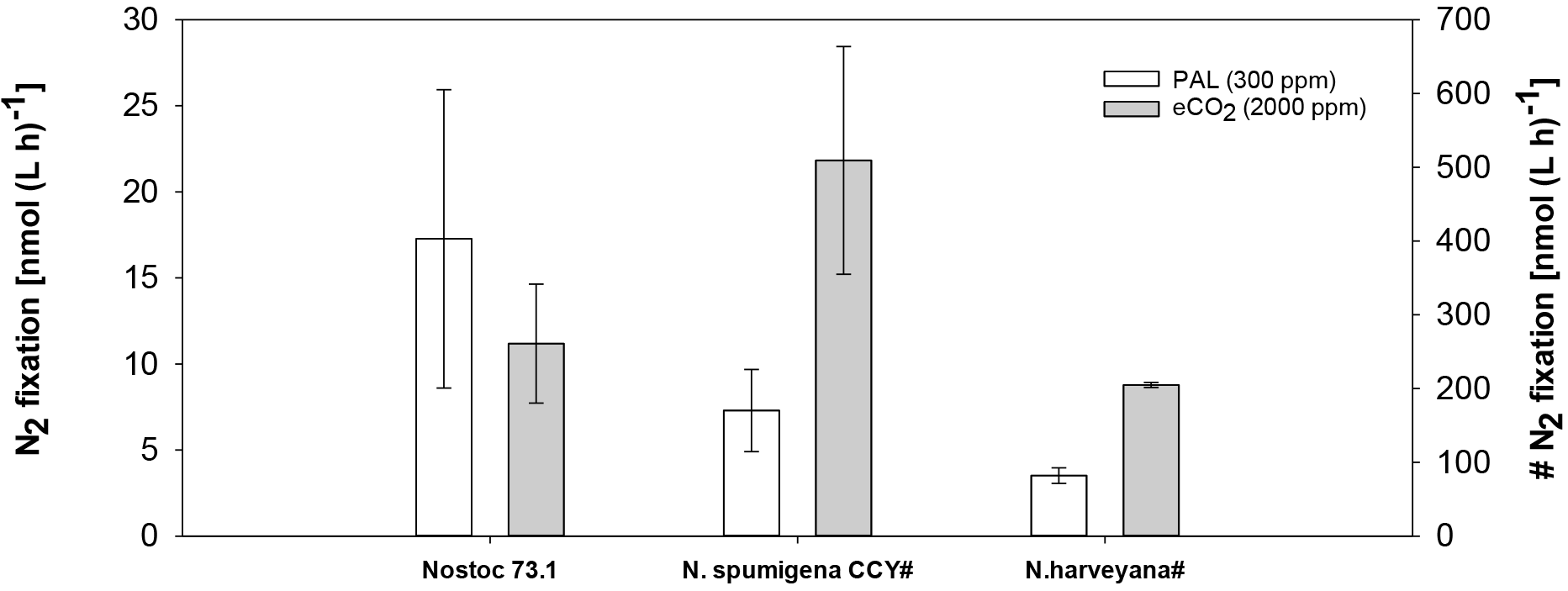
N_2_ fixation rates determined for 3 species grown at PAL (white bars) and eCO_2_ conditions (grey bars). Bars represent mean and standard deviation of two replicates.

### Intracellular toxins

No statistically significant impact of eCO_2_ growth conditions was detected for chlorophyll *a* and volume specific toxin content (Mann-Whitney U test - data not shown, Spearman´s Rho, Suppl. Table 3) in *Nostoc* sp. 73.1, *Nostoc* sp. 65.1 and *N. spumigena* CCY9414. Plotting chlorophyll *a* specific nodularin content as a function of N availability in the media showed a tendency for reduced toxin production in *Nostoc* strains grown in N replete medium at eCO_2_ when compared to PAL controls (Fig. 1 A). This was not the case for *N. spumigena* CCY9414 and *Nostoc* sp. 73.1 cultivated in N-free media, in fact the mean toxin production in *Nostoc* sp. 73.1 appeared greater than in the control (Fig. 1A).

**Figure 4.**
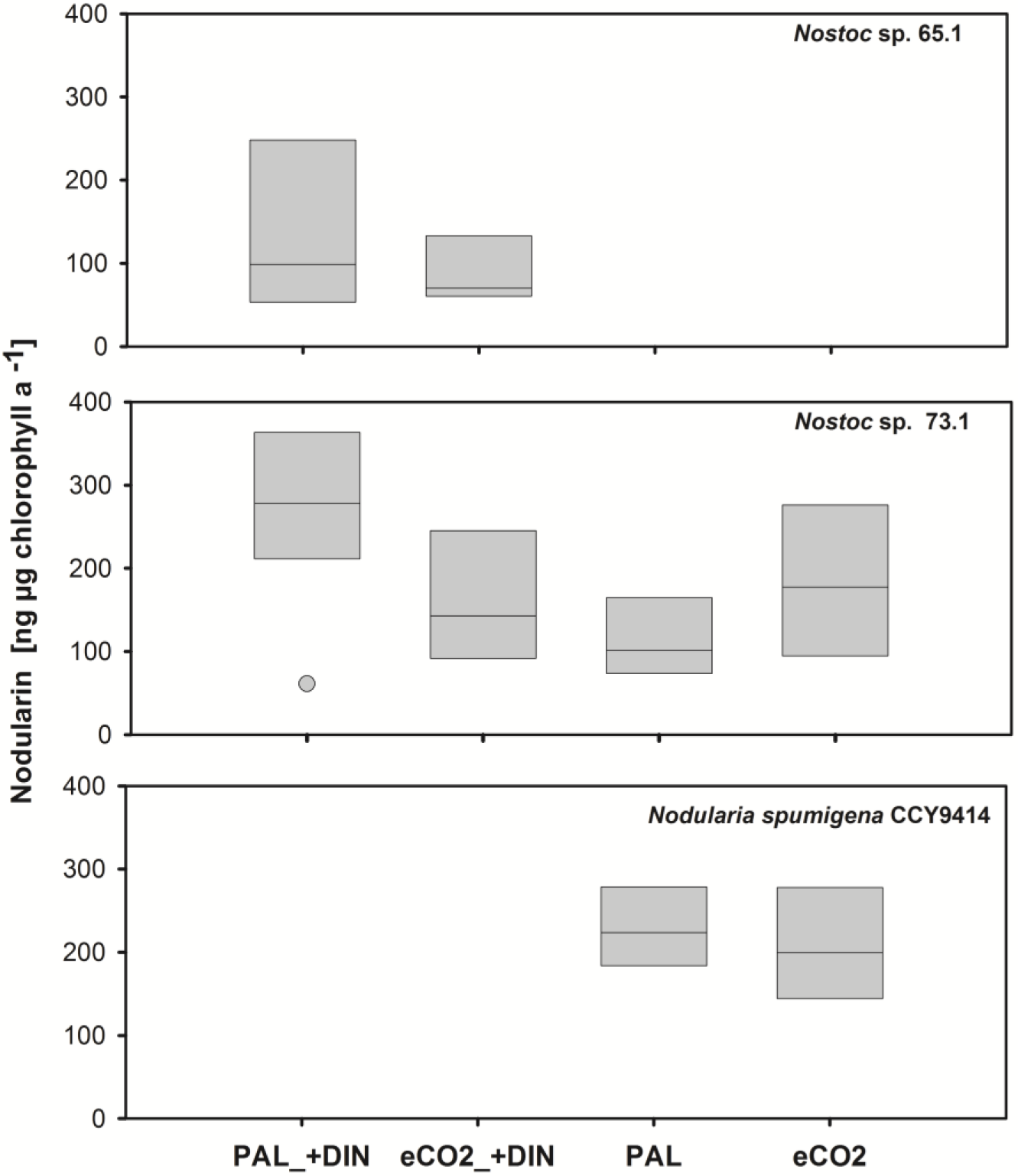
Total soluble cellular nodularin content per μg chlorophyll a as a function of CO_2_ availability of the three toxin producing diazotrophic cyanobacterial strains investigated. *Nodularia spumigena* CCY9414 was grown in DIN-free F/2 medium, *Nostoc* 65.1 in DIN replete BG11 medium and *Nostoc* 73.1 was grown in BG11 and DIN-free BG110 media. Values represent mean and standard deviation after 7 and 14 days incubation (*n* = 4).

We calculated the biomass specific nodularin response ratio (RR) from nodularin content of cells grown at eCO_2_ versus those grown under PAL conditions. Plotting the ln RR against the POC:PON ratio (Fig. 5) highlighted a strain response difference between the terrestrial *Nostoc* nodularin producer and the brackish sweater species *Nodularia spumigena* CCY9414 under N-limiting growth conditions, with the terrestrial *Nostoc* sp. 73.1 exhibiting increased toxin production under eCO_2_, accompanied by an increase in the POC:PON. Particulate matter in *Nostoc* sp. 73.1 grown as a N_2_ fixer displayed significant higher POC:PON ratios compared to the same strain grown in the presence of nitrate. A significant correlation between chlorophyll *a* specific nodularin content and POC:PON ratio was detected (Spearman's rho=0.546, p=0.035, n=15, Suppl. Table 3).

**Figure 5:**
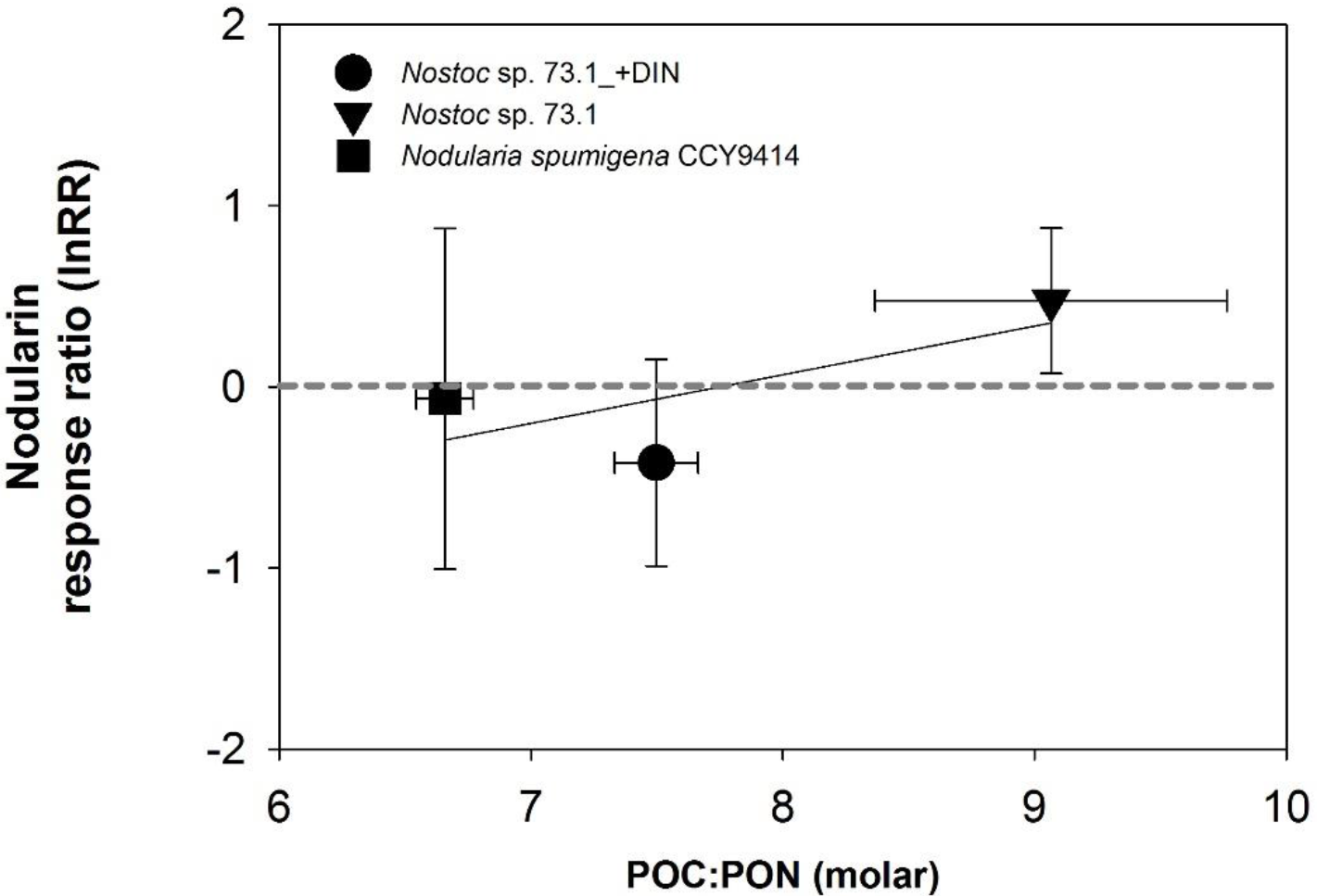
Response ratios of chlorophyll a specific nodularin as a function of particulate organic carbon and nitrogen stoichiometry (POC:PON) in two strains, *Nodularia spumigena* CCY 9414 and *Nostoc* sp. 73.1 with and without DIN in the growth media. The response ratio represents the natural-log transformed contribution of each nodularin variant at eCO_2_ divided by its value at PAL. The horizontal dotted line indicates lack of response (i.e. response ratio = 0), while the bold line indicates linear regressions (*R*^2^ = 0.72, *P* = 0.35). Filled squares represent the response ratio of *Nodularia spumigena* CCY9414, while filled circles and triangles illustrate the response ratio of *Nostoc* sp. 73.1 + DIN and without DIN respectively. Values represent mean ± SD (*n* = 2) after 14 days of incubation.

## Discussion

Rubisco, the enzyme responsible for fixing CO_2_ during oxidative photosynthesis, evolved during the Archean, when atmospheric O_2_ levels were non-existent. The increase in O_2_ in the Earth’s atmosphere, generally accepted to have been brought about by oxidative photosynthesis, is thought to have induced the evolution of the carbon concentrating mechanism to maintain the carboxylase and suppress the oxygenase activity of Rubisco (Rae, 2013). It is well known that cyanobacteria grow faster at elevated levels of CO_2_ (e.g. (Hutchins *et al.*, 2007, Levitan *et al.*, 2007) as they are not reliant on the carbon concentrating mechanisms to maintain high levels of CO_2_ around Rubisco, with some strains, being able to grow at 100% CO_2_ (Thomas *et al.*, 2005). However, the effect of prolonged exposure to eCO_2_ on the NP of cyanobacteria, particularly diazotrophic *Nostoc* and *Nodularia* species, is largely unknown. Our pilot study aimed at elucidating possible changes in NP rates in terrestrial and aquatic diazotrophic cyanobacteria species at near future eCO_2_ levels and additionally measure differences in strain responses with respect to N_2_ fixation and toxin production.

### Net primary production response to eCO_2_

All species and strains tested were able to quickly respond to elevated CO_2_ in the gas exchange cuvette and instantly up-regulate their NP rates (C_high_ vs C_amb_) as illustrated in the short term response graphs (Fig 1A) and the short term response ratios (Suppl. Figure 2A), expect *N. harveyana* SAG44.5 at day 7. Cultures grown at eCO_2_ showed an immediate reduction in NP efficiency when presented with CO_2_ at PAL levels in the cuvette (Fig.1B). The lack of significant reduction in NP rates in cultures grown at eCO_2_ and measured at 380 ppm CO_2_ (T_high_ vs T_amb_) in the cuvette (Suppl. Fig 2B) indicate that the cultures still invested in maintaining their carbon concentrating mechanism while growing at eCO_2_ levels, thereby enabling them to still maintain normal NP rates at PAL of CO_2_. Only *N. harveyana* SAG44.85 showed a significant difference in NP response rates between day 7 and day 14, with an apparently negative response in NP at eCO_2_ on day 7.

Differences of long-term, 14 and 7 day responses (T_high_:C_amb_) are not as pronounced as the short term (T_high_:T_amb_ & C_high_:C_amb_) and most visible in the strains grown in the presence of nitrate (Fig. 2). *Nostoc* strains grown under N limitation did not utilise the availability of eCO_2_ to increase their NP rates as much as those grown without nitrogen limitation, presumably in an attempt to maintain a balanced C:N ratio.

The gas exchange measurements record the CO_2_ taken up by the CO_2_ receptors, NDH-I_3_ and NDH-I_4_, so far identified in all cyanobacterial strains tested (Reviewed in (Price, 2011, Visser *et al.*, 2016). All CO_2_ is converted to HCO_3_^−^ and transported into the CCM where it is converted back to CO_2_ by carbonic anhydrase and is then available to bind to Rubisco. The fact that there was not a significant reduction in NP measured at PAL levels for cultures grown at eCO_2_ (T_amb_) when compared to PAL measurements (C_amb_) of the control cultures (Fig. 1 A & B, Supp. Table 3) suggests that the cyanobacteria maintained their CCM machinery, despite increased CO_2_ availability. Of interest is that media acidification (Suppl. table 1) did not appear to affect NP rates as there were no significant differences between NP rates determined in control or treated cultures at the CO_2_ levels tested (Fig. 1 A&B).

Production of the toxin nodularin by *N. spumigena*, *Nostoc* sp. 73.1 and *Nostoc* sp. 65.1 did not show any response to eCO_2_. On the other hand, nodularin production was correlated to increased POC:PON levels for *Nostoc* sp. 73.1. Here, elevated nodularin was observed in cultures with lower POC:PON ratios in presence of inorganic nitrate in contrast to *Nostoc* sp. 73.1 growing as a N_2_ fixer. Likewise, the cyanobacterium *Microcystis aeruginosa* (non diazotrophic) is known to synthetize nitrogen-rich microcystin variants when exposed to excess N and eCO_2_ (Van de Waal *et al.*, 2009).

### N_2_ fixation response to eCO_2_

The negative yet not significant effect of acidification on N_2_ fixation rates for *Nostoc* sp. 73.1detected in our pilot study should be further investigated. On the other hand, *Nodularia* spp. tested were not inhibited by medium acidification and displayed stimulation at eCO_2_ as published before (Wannicke *et al.*, 2012). This stimulation of N_2_ fixation in *Nodularia* is not consistent among publications with studies reporting negative (Czerny *et al.*, 2009) or non-significant (Karlberg & Wulff, 2013) effect of eCO_2_. Whether this reflects their origin in an acidic aquatic environment in a period of elevated atmospheric CO_2_ levels, remains to be determined.

Freshwater non-diazotrophic cyanobacteria were shown to benefit from increased HCO_3_^−^ in the media only if they carried the low specificity, high flux bicarbonate receptor, BicA (Sandrini *et al.*, 2016, Visser *et al.*, 2016) both published during the execution of this pilot study). As no genetic information is available for the terrestrial strains of *Nostoc* used in this study, nor for *N. harveyana* SAG44.85, we are unable to interpret the biological response to increased HCO_3_^−^, neither within the context of NP nor N_2_ fixation. Excess supply of both C and N is known to increase cellular N:C ratios (Van de Waal *et al.*, 2009). One would therefor expect an increase in N uptake under N-replete conditions or N_2_ fixation in diazotrophs under N-limitation in cultures benefitting from eCO_2_. There appears to be a trend to increased N_2_ fixation rates in *N. spumigena* CCY9414 (Fig. 3) under eCO_2_ growth conditions, with a significant increase recorded for the benthic strain *N. harveyana* SAG44.85. *N. spumigena* CCY9414, for which genomic data is available, carries a gene (NSP_RS09630) with 75% identitity to the *BicA* gene of *M. aeruginosa* PCC7806, thereby suggesting that it can benefit from increased HCO_3_^−^ in the media and increase its NP rates accordingly. Interestingly, the increase in NP did not cause increased respiration (Suppl. Fig. 3), indicating that the cultures were not stressed under eCO_2_ growth conditions and that Rubisco was not running at near substrate saturation (Price, 2011).

### Differences between cyanobacterial species

No matter what Ci is acquired by the cyanobacterium, only CO_2_ converted from HCO_3_^−^ can serve as substrate for Rubisco. Aquatic environments are often severely limited by Ci availability owing to the poor diffusion of CO_2_ in water, and the slow equilibrium between CO_2_ and HCO_3_^−^, especially at neutral pH values. This historical difference in Ci availability is reflected in the responses of terrestrial *Nostoc* species, the bloom forming, surface dwelling *N. spumigena* CCY9414 and the benthic living *N. harveyana* SAG44.85 (Fig. 6). The latter displayed the weakest response towards eCO_2_ in the long term, but the highest increase in short- term response of NP at eCO_2_ in the gas exchange cuvette, suggesting an overall low expression of HCO_3_^−^ receptors. In their natural benthic environment, elevated CO_2_ of up to 3000 |μatm can occur due to high organic matter decomposition and remineralisation (Haynert *et al*., 2012). The capability to swiftly downregulate the CCM is an important prerequisite for attaining high CO_2_ tolerance in this environment.

**Fig. 6.**
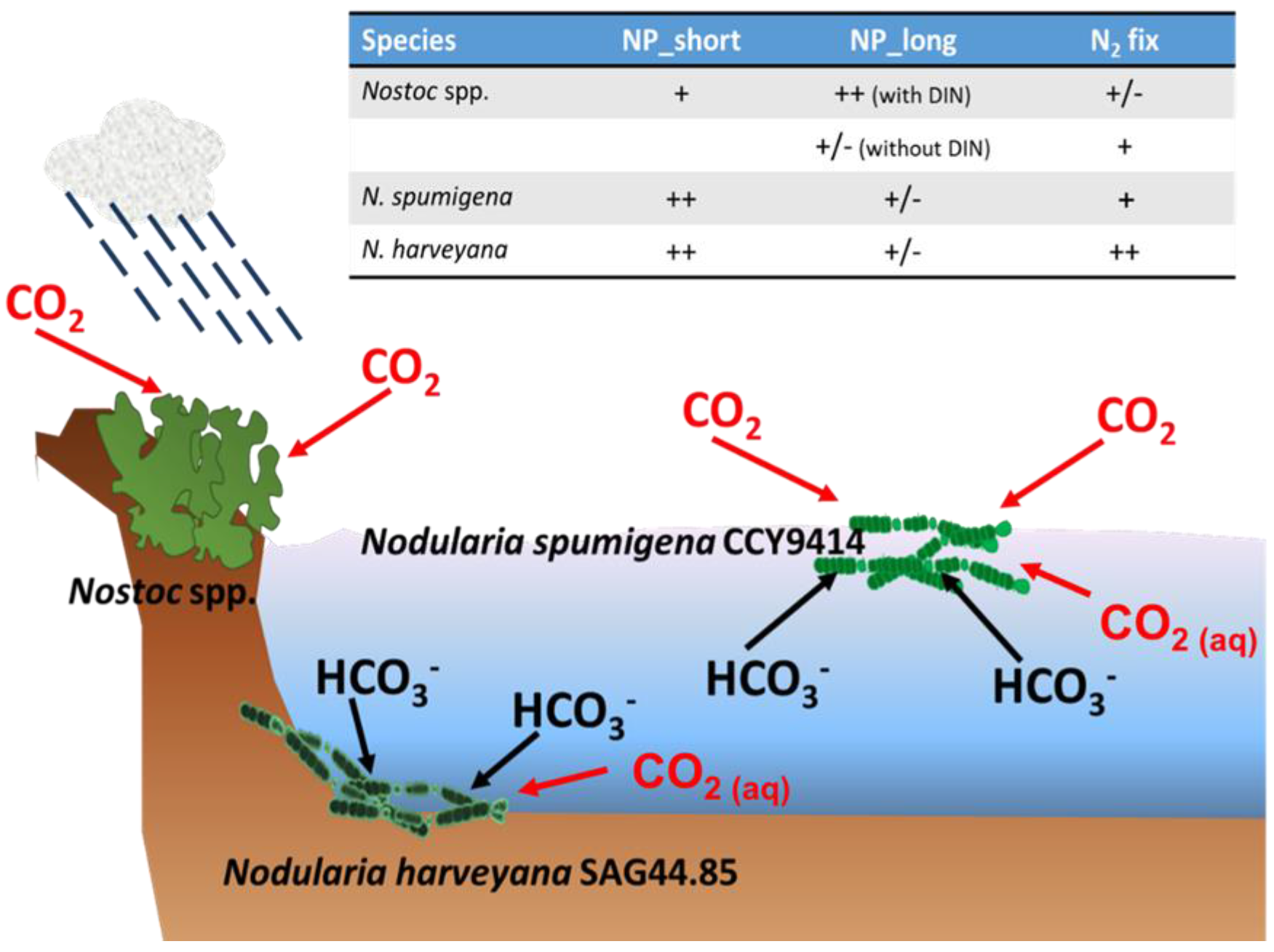
Ecological implications of eCO_2_ on terrestrial, aquatic and benthic cyanobacteria species. Terrestrial cyanobacterial nett photosynthesis (NP) is primarily dependent on the availability of water, while aquatic species are generally restricted by N and P availability. All species can access CO_2_ gas in the atmosphere or dissolved in water. The ability to benefit from increased HCO_3_^−^ availability, the result of increased atmospheric CO_2_ in this study, is largely dependent on whether the species of cyanobacteria expresses the high flux, low specificity sodium dependent BicA carbonate transporter that would allow rapid uptake of HCO_3_^−^ from the water surrounding it. This study only assessed NP based on CO_2_ uptake. Abbreviations: NP: net photosynthesis (short and long-term response), fix: fixation, DIN (dissolved inorganic nitrogen). Mathematic symbols in the table indicate response towards eCO_2_ with +/− representing no response, + slight positive and ++ strong positive response.

*N. spumigena* CCY9414, being an aquatic surface dweller would be exposed to higher levels of Ci both in the water and from the air, thereby allowing for overall increased NP as seen in Fig. 1 when compared to NP of *N. harveyana* SAG44.85. The terrestrial *Nostoc* species used in this study are historically required to be more cautious in their growth as they were isolated from areas prone to long periods of desiccation (Gehringer *et al.*, 2010). While rapidly responding to eCO_2_ with increased NP rates in both the short and long term (Figs. 1 & 2), they did not invest in the highly energy demanding process of N_2_ fixation (Fig. 3) under N limitation as heavily as the aquatic *Nodularia* species studied. The small increase in N_2_ fixation by *Nostoc* sp. 73.1 observed under eCO_2_ in this study may offer an explanation as to the overall reduction in cyanobacterial biomass observed in dryland soilcrusts exposed to eCO_2_ (Steven *et al.*, 2012). This negative effect of exposure to eCO_2_ highlights the complexity of dryland biocrust systems and their response to climate change (Reed *et al.*, 2016). The observed increase in expression of cyanobacterial genes related to oxidative stress (Steven *et al.*, 2012) should be taken into consideration in future studies exposing both terrestrial and aquatic cyanobacterial species to elevated levels of CO_2_.

What is apparent from this pilot study is that there are clear strain specific differences in responding to eCO_2_ levels in strains originating from benthic, pelagic and terrestrial environments, which likely induce changes in the community composition of not only fresh water and marine waters, but also the terrestrial sphere. Future investigations should focus on more clearly elucidating the response of Rubisco to raised atmospheric CO_2_ levels in cyanobacterial strains for which we have complete genomic sequence information. Additionally, the factors governing the large differences in N_2_ fixation rates observed between the *Nodularia* and *Nostoc* strains investigated in the pilot investigations, especially with respect to water acidification, should be further studied. Cyanobacteria have, from a historical perspective, survived numerous environmental challenges during their long lifespan on Earth. This study has shown that all the diazotrophic cyanobacteria investigated are able to immediately increase their NP rates in response to atmospheric eCO_2_, both in the short term and after 14 days exposure. Changes in NP responses to combined elevated CO_2_ and HCO_3_^−^ availability should be assessed in future studies using strains for which we have genetic information pertaining to Ci uptake receptors. The increased PON content and lower POC:PON ratios suggest that increased diazotrophic bloom occurrences will become normality under elevated atmospheric CO_2_ levels up to 2000 ppm, the level investigated in this study. Further studies investigating C sequestration and N_2_ fixation in a wider range of cyanobacterial primary producers from different ecological niches will assist in understanding the ecophysiological responses in fresh- and marine water systems, as well as terrestrial environments.

## Acknowledgments

N.Wannicke thankfully acknowledges the financial support by the Project BIOACID of the German Federal Ministry of Education and Research (BMBF, FKZ 03F0728F). M. Gehringer was partially funded by DFG GE 2558/3-1. We wish to convey our gratitude to B. Büdel, C. Colesie and E. Neuhaus (TU Kaiserslautern) for providing experimental facilities and expertise and E. Dittman, University of Potsdam for toxin analysis.

---

**Supplemental Table 1:** Carbonate chemistry parameter from the cultures after 14 days of experiment either measured or calculated using CO2SYS. Statistical significant differences between the two CO_2_ treatments are indicated by * for p < 0.05 (Student´s t-test, two-tailed). Abbreviations: A_T_ total alkalinity, C_T_ total carbon.

| Species | Growth media | CO<sub>2</sub> treatment | pH measured | A<sub>T</sub> measured<br>[μmol/kg] | C<sub>T</sub> calculated<br>[μmol/kg] | pCO<sub>2</sub> calculated<br>[μatm] |
| --- | --- | --- | --- | --- | --- | --- |
| *Nostoc* | BG110 | PAL (380 ppm) | 7.85± 0.06* | 561± 5 | 504±1 * | 293± 38* |
| *Nodularia* | F/2 | PAL (380 ppm) | 7.85± 0.05* | 455± 1 | 398±3 * | 232± 26* |
| *Nostoc* | BG110 | eCO<sub>2</sub> (2000) | 7.05± 0.01* | 565±6 | 594±5 * | 1987±42* |
| *Nodularia* | F/2 | eCO<sub>2</sub> (2000) | 7.02± 0.02* | 453± 2 | 473±4 * | 1701± 83* |

**Supplemental Table 2:** Statistical results from the two-way repeated measures ANOVA testing the short- term pCO_2_ effects (380 ppm and 1000 ppm) and the interaction with time (sampling day 7 and 14) on net photosynthesis under the two growth conditions PAL (380 ppm) and eCO2 (2000 ppm). Significant differences (p≤0.05) marked in bold. m.s. missing value. n= 4, DF = 1.

| Treatment | Predictor |  | *Nostoc* 65.1<br>+DIN | *Nostoc* 73.1<br>+DIN | *Nostoc* C1.8<br>+DIN | *Nostoc* 73.1 | *Nostoc* C1.8 | *Nodularia*<br>*spumigena* | *Nodularia*<br>*harveyana* |
| --- | --- | --- | --- | --- | --- | --- | --- | --- | --- |
| PAL | CO<sub>2</sub> | F | 42.08 | 13.38 | 4.77 | 1.813 | 37.72 | 3.69 | 6.62 |
|  |  | p | **0.023** | 0.067 | 0.16 | 0.310 | **0.025** | 0.194 | 0.124 |
|  | Time | F | 0.21 | 0.241 | 2.97 | 3.074 | 0.386 | 19.38 | 6.55 |
|  |  | p | 0.692 | 0.672 | 0.232 | 0.222 | 0.598 | **0.048** | 0.125 |
|  | CO<sub>2</sub> x Time | F | 1.00 | 0.091 | 0.026 | 0.739 | 0.084 | 24.75 | 11.92 |
|  |  | p | 0.423 | 0.792 | 0.887 | 0.481 | 0.799 | **0.038** | 0.075 |
| eCO<sub>2</sub> | CO<sub>2</sub> | F | 486.5 | 155.1 | 1.29 | 120.35 | 87.8 | m.s | 0.159 |
|  |  | p | **0.002** | **0.006** | 0.459 | **0.008** | **0.011** |  | 0.799 |
|  | Time | F | 0.381 | 0.303 | 575 | 34.42 | 2.85 | m.s | 6.01 |
|  |  | p | 0.607 | 0.631 | **0.027** | **0.028** | 0.232 |  | 0.134 |
|  | CO<sub>2</sub> x time | F | 1.09 | 12.68 | 31.9 | 25.15 | 1.007 | m.s | 9.46 |
|  |  | p | 0.406 | 0.071 | 0.111 | **0.038** | 0.421 |  | 0.091 |

**Supplemental Table 3:** Spearman´s Rho correlation coefficients among study variables. Significant correlations are denoted with asterisks at significance level \**p* < 0.05 and \*\**p* < 0.01 in bold numbers. Abbreviations are Chla-chlorophyll a, N_2_fix- N_2_ fixation, PAL-present atmospheric levels, PON-particulate organic nitrogen, POC – particulate organic carbon, eCO2 – elevated CO2.

|  |  | Nitrogen | pcCO<sub>2</sub> | Chla | PAL_Dark<br>respiration | PAL_net<br>photosynthesis | eCO<sub>2</sub>_Dark<br>respiration | eCO<sub>2</sub>_net<br>photosynthesis | Total_cellular_<br>nodularin_per Chla | PON | POC | POC:PON | N<sub>2</sub> fixation<br>[per L] | N<sub>2</sub> fixationn<br>[per PON] |
| --- | --- | --- | --- | --- | --- | --- | --- | --- | --- | --- | --- | --- | --- | --- |
| Nitrogen | Spearman´s Rho |  |  | **-,376\*\*** | **-,389\*\*** | .088 | -.214 | -.218 | -.064 | **-,658\*\*** | -.407 | .376 |  |  |
|  | p (two tailed) |  |  | .002 | .002 | .489 | .097 | .092 | .620 | .006 | .118 | .152 | .001 | .001 |
|  | N | 64 | 12 | 64 | 63 | 64 | 61 | 61 | 63 | 16 | 16 | 16 | 16 | 16 |
| pcCO<sub>2</sub> | Spearman´s Rho |  | 1.000 | **-,643\*** | -.510 | .259 | -.294 | -.084 | .441 | .168 | .182 | .217 | -.147 | -.483 |
|  | p (two tailed) |  |  | .024 | .090 | .417 | .354 | .795 | .151 | .602 | .572 | .499 | .649 | .112 |
|  | N | 12 | 12 | 12 | 12 | 12 | 12 | 12 | 12 | 12 | 12 | 12 | 12 | 12 |
| Chla | Spearman´s Rho | **-,376\*\*** | **-,643\*** | 1.000 | **,716\*\*** | **-,700\*\*** | **,714\*\*** | **-,733\*\*** | **-,363\*\*** | .426 | .147 | **-,659\*\*** | **,633\*\*** | **,701\*\*** |
|  | p (two tailed) | .002 | .024 |  | .000 | .000 | .000 | .000 | .003 | .099 | .587 | .006 | .009 | .003 |
|  | N | 64 | 12 | 64 | 63 | 64 | 61 | 61 | 63 | 16 | 16 | 16 | 16 | 16 |
| PAL_Dark respiration | Spearman´s Rho | **-,389\*\*** | -.510 | **,716\*\*** | 1.000 | **-,596\*\*** | **,615\*\*** | **-,614\*\*** | **-,439\*\*** | .482 | .238 | **-,553\*** | **,624\*\*** | **,689\*\*** |
|  | p (two tailed) | .002 | .090 | .000 |  | .000 | .000 | .000 | .000 | .058 | .374 | .026 | .010 | .003 |
|  | N | 63 | 12 | 63 | 63 | 63 | 61 | 61 | 62 | 16 | 16 | 16 | 16 | 16 |
| PAL_net photosynthesis | Spearman´s Rho | .088 | .259 | **-,700\*\*** | **-,596\*\*** | 1.000 | **-,602\*\*** | **,585\*\*** | .101 | **-,694\*\*** | **-,500\*** | **,582\*** | **-,809\*\*** | **-,650\*\*** |
|  | p (two tailed) | .489 | .417 | .000 | .000 |  | .000 | .000 | .430 | .003 | .049 | .018 | .000 | .006 |
|  | N | 64 | 12 | 64 | 63 | 64 | 61 | 61 | 63 | 16 | 16 | 16 | 16 | 16 |
| eCO<sub>2</sub>_Dark respiration | Spearman´s Rho | -.214 | -.294 | **,714\*\*** | **,615\*\*** | **-,602\*\*** | 1.000 | **-,660\*\*** | **-,498\*\*** | .385 | .150 | -.479 | **,580\*** | **,633\*\*** |
|  | p (two tailed) | .097 | .354 | .000 | .000 | .000 |  | .000 | .000 | .141 | .579 | .060 | .019 | .009 |
|  | N | 61 | 12 | 61 | 61 | 61 | 61 | 61 | 60 | 16 | 16 | 16 | 16 | 16 |
| eCO<sub>2</sub>_net photosynthesis | Spearman´s Rho | .218 | -.084 | **-,733\*\*** | **-,614\*\*** | **,585\*\*** | **-,660\*\*** | 1.000 | **,551\*\*** | -.174 | .109 | .474 | -.224 | -.291 |
|  | p (two tailed) | .092 | .795 | .000 | .000 | .000 | .000 |  | .000 | .520 | .688 | .064 | .405 | .274 |
|  | N | 61 | 12 | 61 | 61 | 61 | 61 | 61 | 60 | 16 | 16 | 16 | 16 | 16 |
| Total_cellular_nodularin_per Chla | Spearman´s Rho | -.064 | .441 | **-,363\*\*** | **-,439\*\*** | .101 | **-,498\*\*** | **,551\*\*** | 1.000 | -.052 | .227 | **,546\*** | .014 | -.101 |
|  | p (two tailed) | .620 | .151 | .003 | .000 | .430 | .000 | .000 |  | .853 | .416 | .035 | .959 | .720 |
|  | N | 63 | 12 | 63 | 62 | 63 | 60 | 60 | 63 | 15 | 15 | 15 | 15 | 15 |
| PON | Spearman´s Rho | **-,658\*\*** | .168 | .426 | .482 | **-,694\*\*** | .385 | -.174 | -.052 | 1.000 | **,888\*\*** | -.397 | **,846\*\*** | **,511\*** |
|  | p (two tailed) | .006 | .602 | .099 | .058 | .003 | .141 | .520 | .853 |  | .000 | .128 | .000 | .043 |
|  | N | 16 | 12 | 16 | 16 | 16 | 16 | 16 | 15 | 16 | 16 | 16 | 16 | 16 |
| POC | Spearman´s Rho | -.407 | .182 | .147 | .238 | **-,500\*** | .150 | .109 | .227 | **,888\*\*** | 1.000 | 0.000 | **,667\*\*** | .246 |
|  | p (two tailed) | .118 | .572 | .587 | .374 | .049 | .579 | .688 | .416 | .000 |  | 1.000 | .005 | .359 |
|  | N | 16 | 12 | 16 | 16 | 16 | 16 | 16 | 15 | 16 | 16 | 16 | 16 | 16 |
| POC:PON | Spearman´s Rho | .376 | .217 | **-,659\*\*** | **-,553\*** | **,582\*** | -.479 | .474 | **,546\*** | -.397 | 0.000 | 1.000 | **-,525\*** | **-,570\*** |
|  | p (two tailed) | .152 | .499 | .006 | .026 | .018 | .060 | .064 | .035 | .128 | 1.000 |  | .037 | .021 |
|  | N | 16 | 12 | 16 | 16 | 16 | 16 | 16 | 15 | 16 | 16 | 16 | 16 | 16 |
| N<sub>2</sub> fixation [per L] | Spearman´s Rho | **-,752\*\*** | -.147 | **,633\*\*** | **,624\*\*** | **-,809\*\*** | **,580\*** | -.224 | .014 | **,846\*\*** | **,667\*\*** | **-,525\*** | 1.000 | **,847\*\*** |
|  | p (two tailed) | .001 | .649 | .009 | .010 | .000 | .019 | .405 | .959 | .000 | .005 | .037 |  | .000 |
|  | N | 16 | 12 | 16 | 16 | 16 | 16 | 16 | 15 | 16 | 16 | 16 | 16 | 16 |
| N<sub>2</sub> fixationn [per PON] | Spearman´s Rho | **-,752\*\*** | -.483 | **,701\*\*** | **,689\*\*** | **-,650\*\*** | **,633\*\*** | -.291 | -.101 | **,511\*** | .246 | **-,570\*** | **,847\*\*** | 1.000 |
|  | p (two tailed) | .001 | .112 | .003 | .003 | .006 | .009 | .274 | .720 | .043 | .359 | .021 | .000 |  |
|  | N | 16 | 12 | 16 | 16 | 16 | 16 | 16 | 15 | 16 | 16 | 16 | 16 | 16 |

**Supplemental Figure 1:**
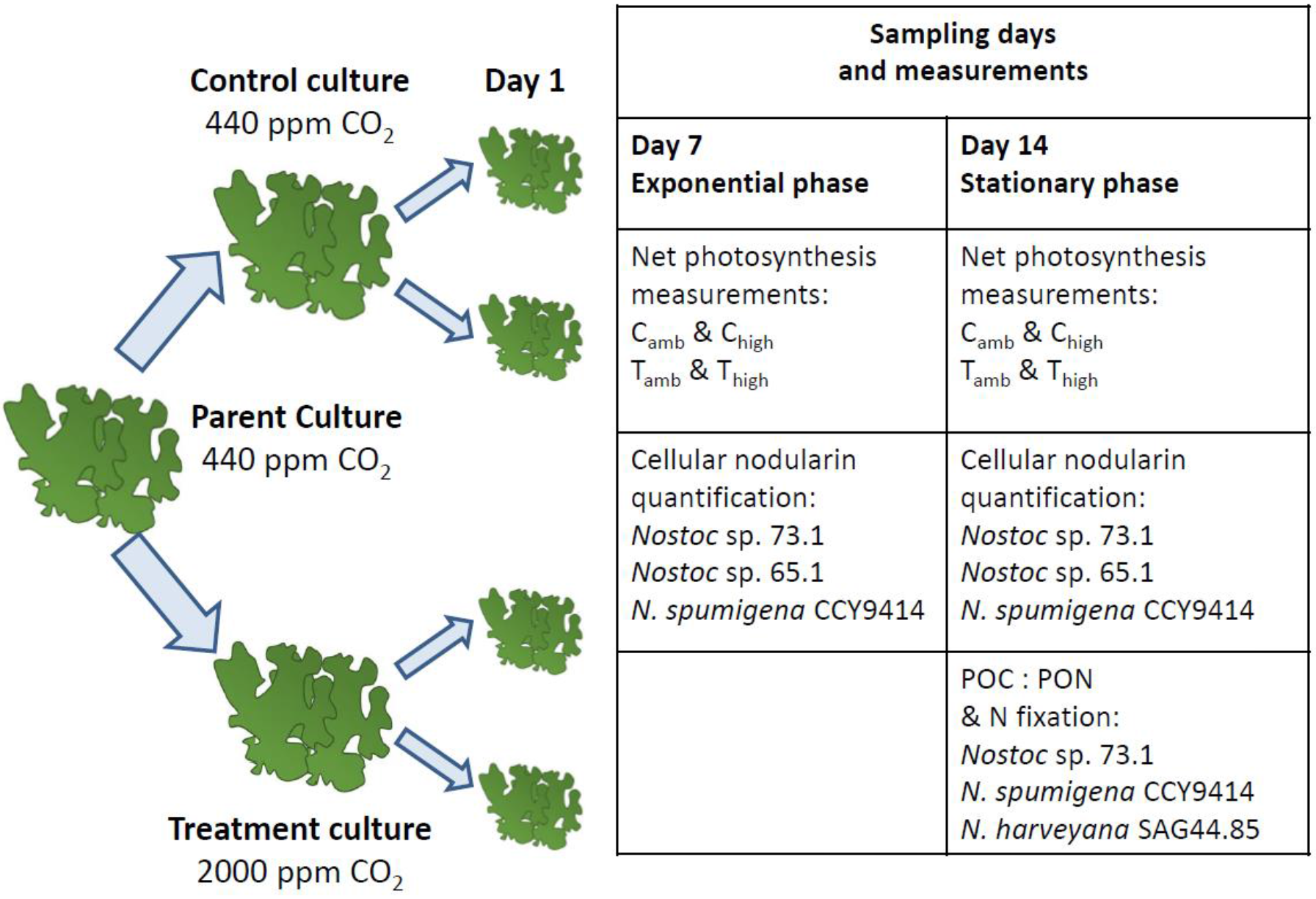
Diagram illustrating the experimental flow. Parent cultures grown under PAL conditions were divided into two batches 14 days prior to the start of the experiment to allow for acclimation to the elevated CO2 levels. These cultures were used as inocula for the experimental cultures and processed as indicated and described in the main text.

**Supplemental Figure 2:**
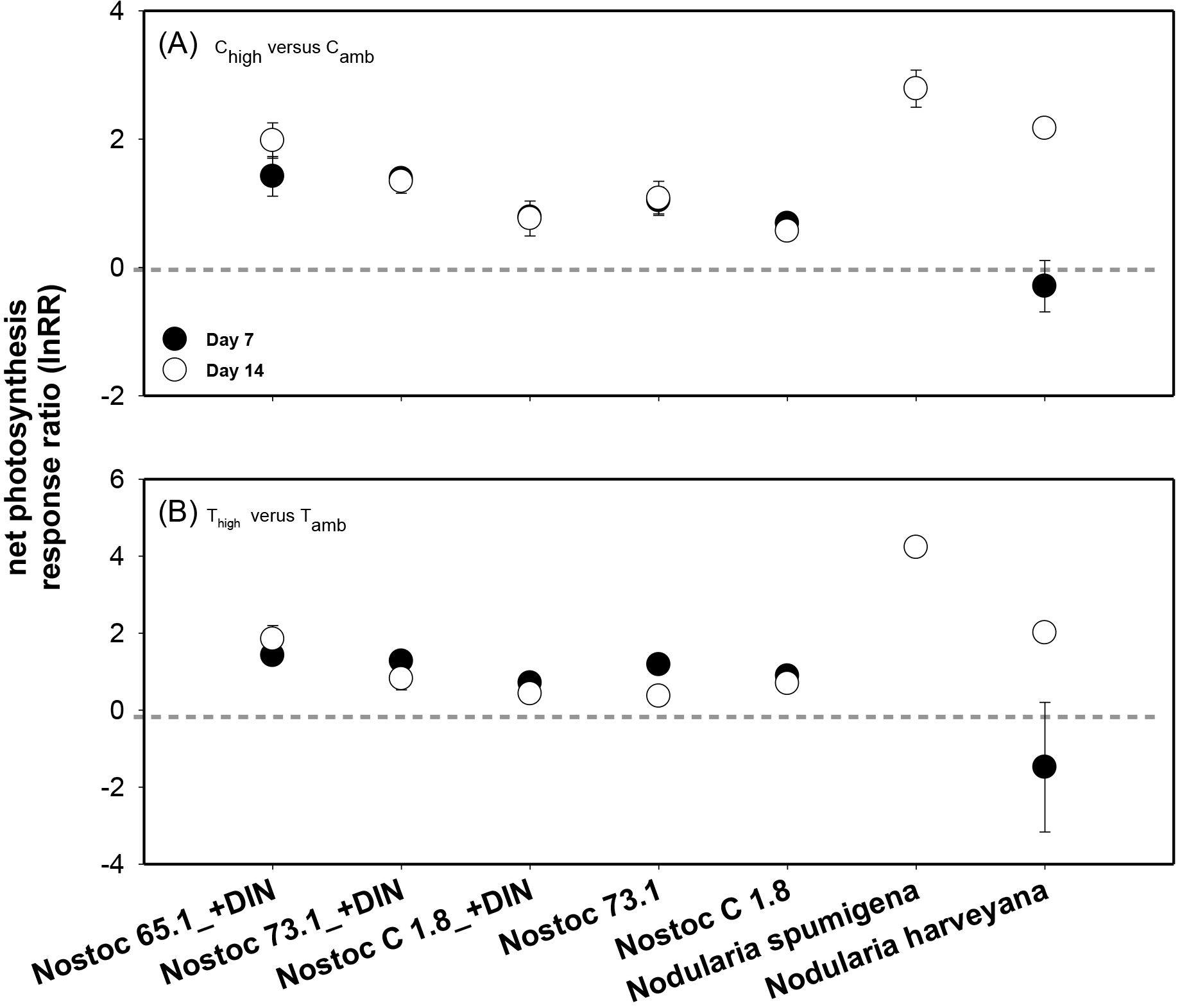
Short term response of net photosynthesis for the different species tested. The lnRR (natural-log transformed) was calculated from the quotient of the net photosynthesis (NP) rate of C_high_ by that of C_amb_ (A). In (B), the lnRR of T_high_ divided by the NP rates of T_amb_ is plotted. The lnRR data is presented calculated from the means and standard deviations of days 7 and 14, n=2 respectively. The horizontal grey line indicates a lack of response to the eCO_2_ treatment (i.e. lnRR = 0). C_high_ readings for day 7 for *N. spumigena* PCC9414 were erratic and not included in the analysis.

**Supplemental Figure 3:**
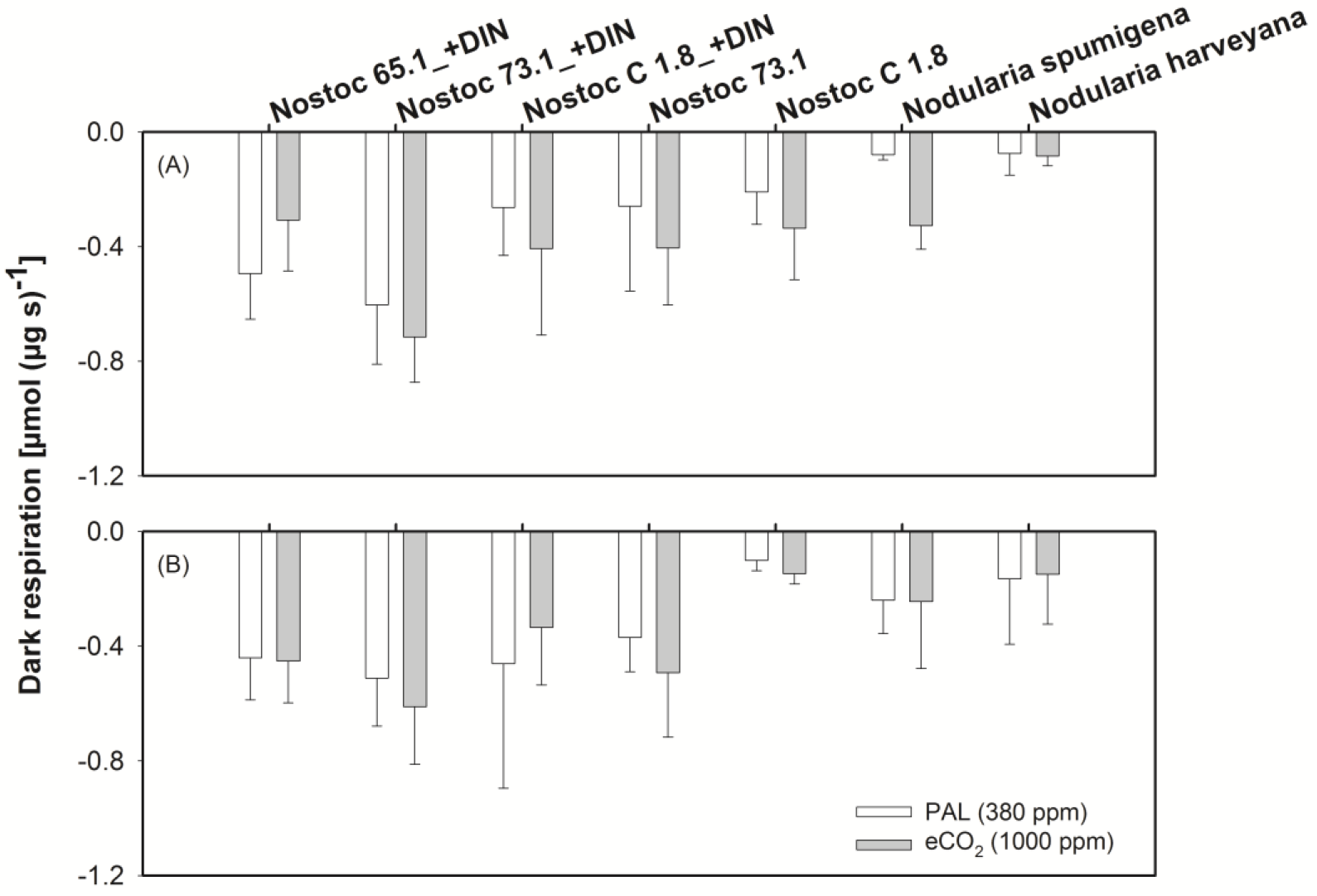
Short term acclimation responses of dark respiration rates of cultures (A) grown at PAL and (B) grown at eCO_2_. NP measured under PAL (white bars) and eCO_2_ (grey bars). Bars represent the mean and standard deviation of n=4 (days 7+ 14).

## References

Beversdorf LJ, Miller TR & McMahon KD (2013) The Role of Nitrogen Fixation in Cyanobacterial Bloom Toxicity in a Temperate, Eutrophic Lake. Plos One 8.

Bhargava S, Chouhan S, Kaithwas V & Maithil R (2013) Carbon dioxide regulation of autotrophy and diazotrophy in the nitrogen-fixing cyanobacterium Nostoc muscorum. Ecotox Environ Safe 98: 345–351.

Cardona T, Murray JW & Rutherford AW (2015) Origin and Evolution of Water Oxidation before the Last Common Ancestor of the Cyanobacteria. Molecular Biology and Evolution 32: 1310–1328.

Czerny J, Barcelos e Ramos J & Riebesell U (2009) Influence of elevated CO2 concentrations on cell division and nitrogen fixation rates in the bloom-forming cyanobacterium Nodularia spumigena. Biogeosciences (BG) 6: 1865–1875.

Downing TG, Sember CS, Gehringer MM & Leukes W (2005) Medium N : P ratios and specific growth rate comodulate microcystin and protein content in Microcystis aeruginosa PCC7806 and M-Aeruginosa UV027. Microbial Ecol 49: 468–473.

Eichner M, Rost B & Kranz SA (2014) Diversity of ocean acidification effects on marine N-2 fixers. J Exp Mar Biol Ecol 457: 199–207.

El-Shehawy R, Gorokhova E, Fernandez-Pinas F & del Campo FF (2012) Global warming and hepatotoxin production by cyanobacteria: What can we learn from experiments? Water Research 46: 1420–1429.

Falkowski P & Raven J (2007) Photosynthesis and primary production in nature. Aquatic photosynthesis 2nd ed Princeton University Press, Princeton.

Gehringer MM (2004) Microcystin-LR and okadaic acid-induced cellular effects: A dualistic response. FEBS Letters 557: 1–8.

Gehringer MM & Wannicke N (2014) Climate change and regulation of hepatotoxin production in Cyanobacteria. Fems Microbiol Ecol 88: 1–25.

Gehringer MM, Pengelly JJL, Cuddy WS, Fieker C, Forster PI & Neilan BA (2010) Host selection of symbiotic cyanobacteria in 31 species of the Australian cycad genus: Macrozamia (Zamiaceae). Molecular Plant-Microbe Interactions 23: 811–812.

Gehringer MM, Adler L, Roberts AA, Moffitt MC, Mihali TK, Mills TJ, Fieker C & Neilan BA (2012) Nodularin, a cyanobacterial toxin, is synthesized in planta by symbiotic Nostoc sp. The ISME journal 6: 1834–1847.

Haynert K, Schönfeld J, Polovodova-Asteman I & Thomsen J (2012) The benthic foraminiferal community in a naturally CO2-rich coastal habitat in the southwestern Baltic Sea. Biogeosciences (BG) 9: 4421–4440

Horst GP, Sarnelle O, White JD, Hamilton SK, Kaul RB & Bressie JD (2014) Nitrogen availability increases the toxin quota of a harmful cyanobacterium, Microcystis aeruginosa. Water Research 54: 188–198.

Hutchins DA, Fu FX, Zhang Y, Warner ME, Feng Y, Portune K, Bernhardt PW & Mulholland MR (2007) CO2 control of Trichodesmium N-2 fixation, photosynthesis, growth rates, and elemental ratios: Implications for past, present, and future ocean biogeochemistry. Limnol Oceanogr 52: 1293–1304.

Hutchins DA, Fu FX, Zhang Y, Warner ME, Feng Y, Portune K, Bernhardt PW & Mulholland MR (2007) CO2 control of Trichodesmium N-2 fixation, photosynthesis, growth rates, and elemental ratios: Implications for past, present, and future ocean biogeochemistry. Limnology and Oceanography 52: 1293–1304.

Karlberg M & Wulff A (2013) Impact of temperature and species interaction on filamentous cyanobacteria may be more important than salinity and increased pCO2 levels. Mar Biol 160: 2063–2072.

Kleinteich J, Wood SA, Küpper FC, Camacho A, Quesada A, Frickey T & Dietrich DR (2012) Temperature-related changes in polar cyanobacterial mat diversity and toxin production. Nature Climate Change 2: 356–360.

Kranz SA, Eichner M & Rost B (2011) Interactions between CCM and N-2 fixation in Trichodesmium. Photosynth Res 109: 73–84.

Lane RW, Menon M, McQuaid JB, Adams DG, Thomas AD, Hoon SR & Dougill AJ (2013) Laboratory analysis of the effects of elevated atmospheric carbon dioxide on respiration in biological soil crusts. J Arid Environ 98: 52–59.

Levitan O, Rosenberg G, Setlik I, Setlikova E, Grigel J, Klepetar J, Prasil O & Berman-Frank I (2007) Elevated CO2 enhances nitrogen fixation and growth in the marine cyanobacterium Trichodesmium. Global Change Biol 13: 531–538.

Lewis CE, Causton DR, Peratoner G, Cairns AJ & Foyer CH (1998) Acclimation of Lolium temulentum to growth at elevated CO2. Photosynthesis: Mechanisms and Effects, Vols I-V 4035–4038.

Liu J, Van Oosterhout E, Faassen EJ, Lurling M, Helmsing NR & Van de Waal DB (2016) Elevated pCO(2) causes a shift towards more toxic microcystin variants in nitrogen-limited Microcystis aeruginosa. Fems Microbiol Ecol 92, fiv159.

Lyons TW, Reinhard CT & Planavsky NJ (2014) The rise of oxygen in Earth's early ocean and atmosphere. Nature 506: 307–315.

Meeks JC & Castenholz RW (1971) Growth and Photosynthesis in an Extreme Thermophile, Synechococcus-Lividus (Cyanophyta). Arch Mikrobiol 78: 25–41.

Mohr W, Grosskopf T, Wallace DWR & LaRoche J (2010) Methodological Underestimation of Oceanic Nitrogen Fixation Rates. Plos One 5.

Montoya JP, Voss M, Kahler P & Capone DG (1996) A simple, high-precision, high-sensitivity tracer assay for N-2 fixation. Appl Environ Microb 62: 986–993.

Neilan BA, Pearson LA, Muenchhoff J, Moffitt MC & Dittmann E (2013) Environmental conditions that influence toxin biosynthesis in cyanobacteria. Environ Microbiol 15: 1239–1253.

Paerl HW & Huisman J (2009) Climate change: a catalyst for global expansion of harmful cyanobacterial blooms. Environ Microbiol Rep 1: 27–37.

Paerl HW & Huisman J (2009) Climate change: a catalyst for global expansion of harmful cyanobacterial blooms. Env Microbiol Rep 1: 27–37.

Posch T, Köster O, Salcher MM & Pernthaler J (2012) Harmful filamentous cyanobacteria favoured by reduced water turnover with lake warming. Nature Climate Change 2: 809–813.

Price GD (2011) Inorganic carbon transporters of the cyanobacterial CO2 concentrating mechanism. Photosynth Res 109: 47–57.

Reed SC, Maestre FT, Ochoa-Hueso R, Kuske CR, Darrouzet-Nardi A, Oliver M, Darby B, Sancho LG, Sinsabaugh RL & Belnap J (2016) Biocrusts in the Context of Global Change. Biological Soil Crusts: An Organizing Principle in Drylands, (Weber B, Büdel B & Belnap J, eds.), pp. 451–476. Springer International Publishing, Cham.

Sanchez-Baracaldo P (2015) Origin of marine planktonic cyanobacteria. Sci Rep-Uk 5.

Sandrini G, Jakupovic D, Matthijs HCP & Huisman J (2015) Strains of the Harmful Cyanobacterium Microcystis aeruginosa Differ in Gene Expression and Activity of Inorganic Carbon Uptake Systems at Elevated CO2 Levels. Appl Environ Microb 81: 7730–7739.

Sandrini G, Ji X, Verspagen JMH, Tann RP, Slot PC, Luimstra VM, Schuurmans JM, Matthijs HCP & Huisman J (2016) Rapid adaptation of harmful cyanobacteria to rising CO2. P Natl Acad Sci USA 113:9315–9320.

Sevilla E, Martin-Luna B, Bes MT, Fillat MF & Peleato ML (2012) An active photosynthetic electron transfer chain required for mcyD transcription and microcystin synthesis in Microcystis aeruginosa PCC7806. Ecotoxicology 21: 811–819.

Shi D, Kranz SA, Kim JM & Morel FMM (2012) Ocean acidification slows nitrogen fixation and growth in the dominant diazotroph Trichodesmium under low-iron conditions. P Natl Acad Sci USA 109: E3094–E3100.

Steven B, Gallegos-Graves L, Yeager CM, Belnap J, Evans RD & Kuske CR (2012) Dryland biological soil crust cyanobacteria show unexpected decreases in abundance under long-term elevated CO2. Environ Microbiol 14: 3247–3258.

Thomas DJ, Sullivan SL, Price AL & Zimmerman SM (2005) Common freshwater cyanobacteria grow in 100% CO2. Astrobiology 5: 66–74.

Van de Waal DB, Verspagen JMH, Lurling M, Van Donk E, Visser PM & Huisman J (2009) The ecological stoichiometry of toxins produced by harmful cyanobacteria: an experimental test of the carbon-nutrient balance hypothesis. Ecology Letters 12: 1326–1335.

Van de Waal DB, Verspagen JM, Lürling M, Van Donk E, Visser PM & Huisman J (2009) The ecological stoichiometry of toxins produced by harmful cyanobacteria: an experimental test of the carbon-nutrient balance hypothesis. Ecology letters 12: 1326–1335.

Van de Waal DB, Verspagen JMH, Finke JF, et al. (2011) Reversal in competitive dominance of a toxic versus non-toxic cyanobacterium in response to rising CO2. Isme J 5: 1438–1450.

Verschoor AM, Van Dijk MA, Huisman J & Van Donk E (2013) Elevated CO2 concentrations affect the elemental stoichiometry and species composition of an experimental phytoplankton community. Freshwater Biol 58: 597–611.

Visser PM, Verspagen JMH, Sandrini G, Stal LJ, Matthijs HCP, Davis TW, Paerl HW & Huisman J (2016) How rising CO2 and global warming may stimulate harmful cyanobacterial blooms. Harmful Algae 54: 145–159.

Voss B, Bolhuis H, Fewer DP, et al. (2013) Insights into the Physiology and Ecology of the Brackish-Water-Adapted Cyanobacterium Nodularia spumigena CCY9414 Based on a Genome-Transcriptome Analysis. Plos One 8.

Wannicke N, Endres S, Engel A, Grossart HP, Nausch M, Unger J & Voss M (2012) Response of Nodularia spumigena to pCO(2) - Part 1: Growth, production and nitrogen cycling. Biogeosciences 9: 2973–2988.

Wannicke N, Endres S, Engel A, Grossart HP, Nausch M, Unger J & Voss M (2012) Response of Nodularia spumigena to pCO2 - Part 1: Growth, production and nitrogen cycling. Biogeosciences 9: 2973–2988.

Weiss MC, Sousa FL, Mrnjavac N, Neukirchen S, Roettger M, Nelson-Sathi S & Martin WF (2016) The physiology and habitat of the last universal common ancestor. Nat Microbiol 1, article 16116.

